# Systemic activation coordinates the heat shock response of the insulin/IGF-1 pathway in *Caenorhabditis elegans*

**DOI:** 10.1101/131375

**Authors:** Ronen B Kopito, Kathie Watkins, Erel Levine

## Abstract

Exposure to high temperatures has an adverse effect on cellular processes and results in activation of the cellular heat shock response (HSR), a highly conserved program of inducible genes to maintain protein homeostasis^1^. The insulin/IGF-1 signaling (IIS) pathway, which has diverse roles from metabolism to stress response and longevity, is activated as part of the HSR^2–4^. Recent evidence suggest that the IIS pathway is able to affect proteostasis non-autonomously^5,6^, yet it is not known if it is activated autonomously in stressed cells or systemically as part of an organismic program. In *Caenorhabditis elegans*, the single forkhead box O (FOXO) homologue DAF-16 functions as the major target of the IIS pathway^7^ and, together with the heat-shock factor HSF-1, induce the expression of small heat shock proteins in response to heat shock^8–10,3^. Here we use a novel microfluidic device that allows precise control of the spatiotemporal temperature profile to show that cellular activation of DAF-16 integrates local temperature sensation with systemic signals. We demonstrate that DAF-16 activation in head sensory neurons is essential for DAF-16 activation in other tissues, but show that no known thermosensory neuron is individually required. Our findings demonstrate that systemic and cell-autonomous aspects of stress response act together to facilitate a coordinated cellular response at the organismic level.

---

Genetic evidence suggest potential role of IIS in cell non-autonomous heat-shock response^5,6,11^ and raise the question whether the pathway is activated in stressed cells to initiate non-autonomous response or is activated by systemic signals. To address this question we sought to characterize the spatiotemporal dynamics of IIS activation following heat-shock by performing longitudinal time-lapse imaging of transgenic worms that express DAF-16::GFP (hereafter “DAF-16” for short) from a single chromosomal copy^12^. Activation of the IIS pathway results in translocation of DAF-16 to cell nuclei to modulate the expression of downstream genes and in accumulation of nuclear fluorescence, observable at a single cell level (**Fig. 1**)^2^. Long-term longitudinal imaging and precise temperature control were made possible by a specialized microfluidic device^13^. In this device, worms are individually confined to optimized chambers, and a heat reservoir placed immediately above the chambers is used to control their temperature (**Fig. 1a**). We have previously established that under normal conditions, worms in the device exhibit normal physiological cues and are not stressed^13^.

**Figure 1.**
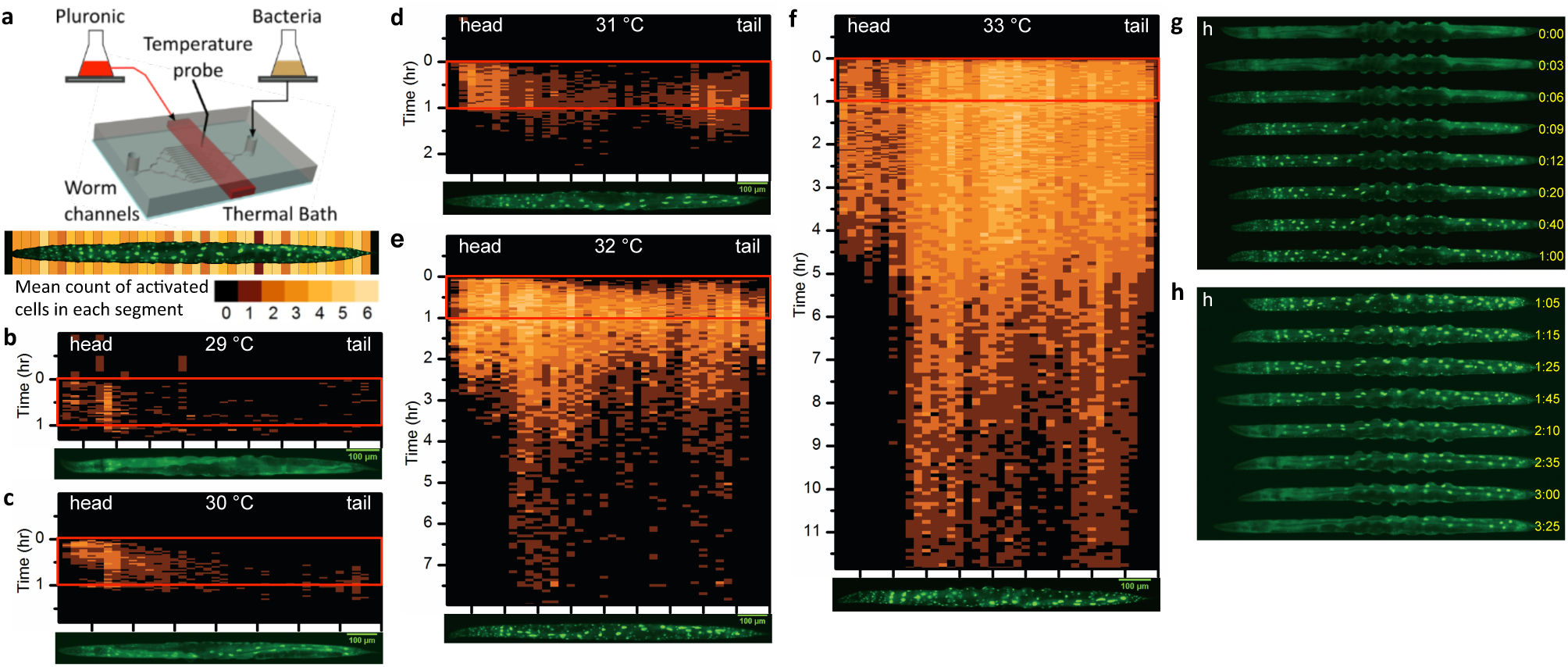
DAF-16 heat shock response is temperature dependent. **a,** Experimental set-up and data analysis. Worms maintained in chambers of a microfluidic device were constantly supplied with food. Temperature was controlled by flowing fluid (Pluronic solution) through a second layer channel (in red) and probed by a thermocouple in close proximity to the worms. For analysis, at each time point nuclei showing DAF-16::GFP fluorescence accumulation were automatically detected by a heuristic image analysis procedure. Worms were segmented into 40 segments along the anterior-posterior (AP) axis, and the number of detected nuclei in each segment was counted, averaged over multiple worms (*n*>25 per condition), and presented by a color code. **b-f,** Dynamics of Daf-16 heat shock response at different temperatures (29°C - 33°C respectively). Vertical axis: time (hours), horizontal axis: worm AP axis, head on the left and tail on the right. In all experiments, one-hour heat shock is applied at time zero. A representative worm at the end of the heat shock is presented for each temperature. **g-h,** Both activation (g) and recovery (f) progress from head to tail.

In all experiments, synchronized populations of worms were raised at 24**°**C to young adulthood on standard solid-media plates, loaded into the device, and allowed a two-hour acclimation period. Under these conditions, worms displayed a diffuse GFP fluorescence throughout their body, indicating that DAF-16 was present both in and out of cell nuclei. A one-hour heat-shock resulted in accumulation of fluorescence in many nuclei, a hallmark of DAF-16 activation^2^. Remarkably, we found that in the range 29-33**°**C the spatial pattern of nuclear accumulation was strongly temperature dependent (**Fig. 1b-f and Movies 1-3**) and highly reproducible between worms (**Extended Data Fig. 1**). In particular we found that the minimal required temperature for DAF-16 translocation was different in different tissues: while head neurons and intestinal cells showed DAF-16 activation at all temperatures in this range, cells in other tissues showed nuclear accumulation only at higher temperatures (**Extended Data Fig. 2**). Moreover, the intensity of nuclear accumulation (that is, the ratio between fluorescence in and out of cell nuclei) increased significantly with temperature (**Extended Data Fig. 3**). These data suggest the existence of tissue-dependent temperature thresholds for DAF-16 activation.

If DAF-16 is activated non-autonomously then the temporal order of activation in individual cells could be sequential and reproducible between worms. This was indeed the case at all the measured temperatures. At temperatures below 33**°**C the precise timing of nuclear accumulation in each tissue was temperature dependent, but their temporal order remained invariant. Importantly, head neurons were always the first to show nuclear DAF-16 accumulation, followed by intestinal cells. Moreover, within each tissue we observed clear progression from head to tail (**Fig. 1b-e,g** and **Extended Data Fig. 2**). In contrast, heat shock of 33**°**C and above resulted in immediate nuclear accumulation throughout the worm body (**Fig. 1f**). Intriguingly, recovery from heat shock was characterized by a similar robust dynamical pattern, displaying an invariant order among tissues and a consistent progression from head to tail (**Fig. 1b-f** and **Fig. 1h**). These data suggest a coordinated systemic activation and deactivation of the IIS pathway across the worm in response to heat shock.

To differentiate between cell-autonomous and non-autonomous components of DAF-16 heat response, we asked if activation by heat shock in one part of the worm is sufficient to induce activation in an unstressed part. To address this question we split the water bath in the microfluidic device into two compartments such that the anterior and posterior parts of the worm could be held at different temperatures (**Fig 2a and Movie 4**). Exposing the anterior part of the worms to a heat shock of either 31 or 33**°**C resulted in activation of cells only in their anterior part (**Fig 2b**). Conversely, worms that experienced a heat shock only at the posterior part of the body showed two types of response. At 31**°**C, about 60% of the observed worms (*n*=53) showed no response at all. The remaining worms, and all worms at 33**°**C, responded first in head neurons and later throughout the heated posterior region (**Fig 2c**). Together we observe that activation of DAF-16 either occurs first in head neurons and then exclusively in the heated part of the worm, or does not occur at all. Thus DAF-16 activation in heated cell coincides with DAF-16 activation in some head neurons^11^.

**Figure 2.**
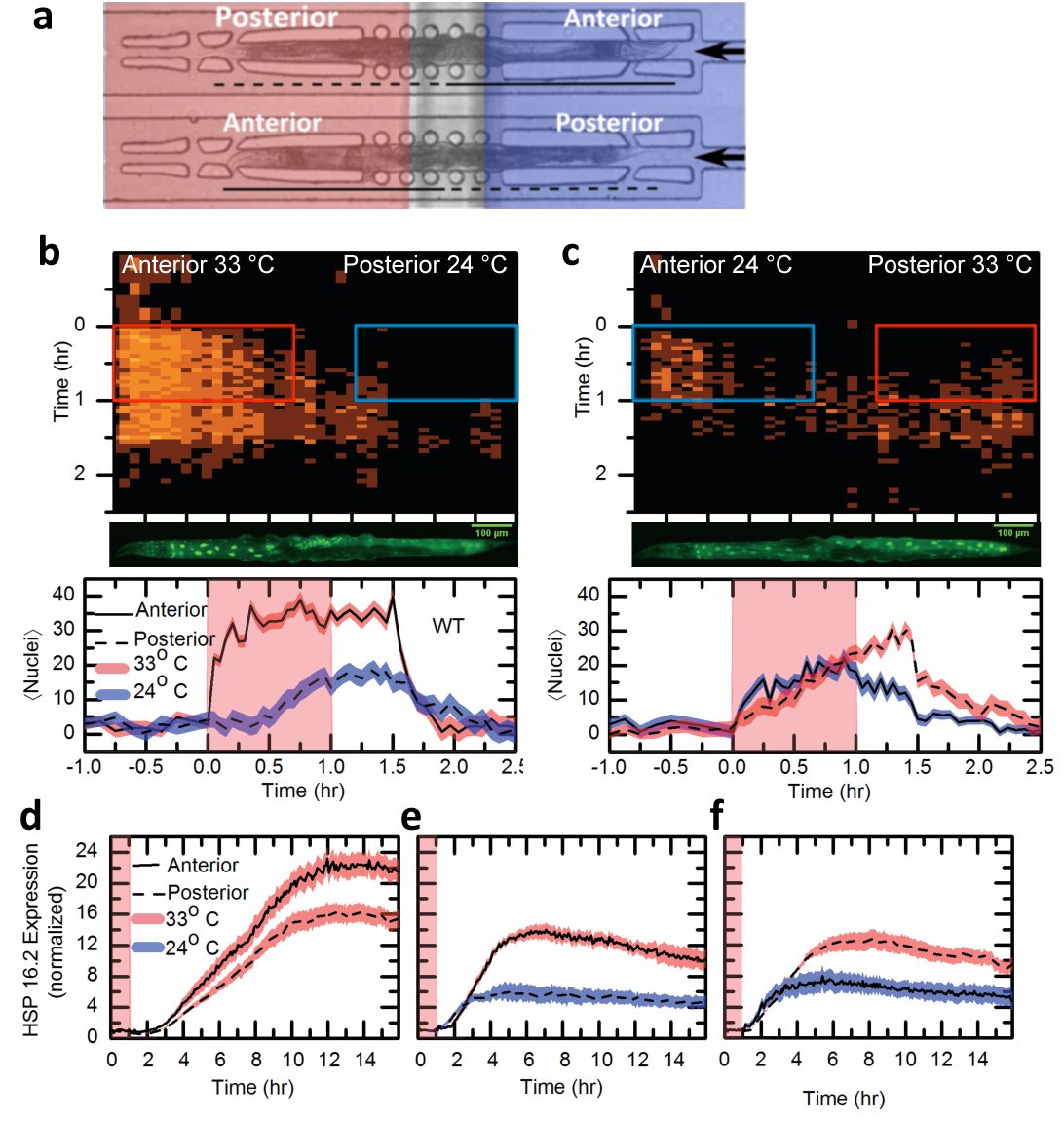
DAF-16 heat shock response to a spatial temperature step. **a,** Two channels separated by a PDMS membrane (200 um) were positioned immediately under the worm chambers in the microfluidic device. Since orientation of the worms in the device was random, about half of the worms are heat shocked at the anterior part and the other half at the posterior. **b-c**, DAF-16 activation dynamics represented as in Fig.1 for anterior heat-shock (b) and posterior heat-shock (c). Blue and red boxes signify the regions directly underneath the 24°C and 33°C heat-baths during heat shock. Middle, a representative worm at the end of the heat shock. Bottom, average number of nuclei with localized fluorescence at the anterior (solid line) and posterior (dashed line) parts of the worm (*n*>20). **d-f,** Normalized mean fluorescence for HSP 16.2::GFP reporter after one hour of heat shock, full body (d), anterior part (e) or posterior part (f).

Intriguingly, the activation pattern in the heated part of the animal was much weaker compared with worms that were exposed to a uniform heat shock: the number of activated cells was smaller, the ratio between nuclear and cytoplasmic fluorescence was lower, and recovery started earlier (only 30 minutes after return from 33**°**C to ambient temperature, cf. 3 hours after a uniform heat shock) and progressed faster (completed in less than 20 minutes rather than more than 5 hours). This suggests that the overall level of heat shock across the animal body is integrated and processed by the neural network to produce a coordinated systemic activation of DAF-16 across the animal body.

To show that DAF-16 translocation pattern correlates with activation of heat-shock response we followed the expression of the small heat-shock protein HSP-16, whose induction under heat-shock requires the heat shock factor HSF-1 as well as DAF-16^10^. Following a uniform heat shock of 33**°**C, a GFP reporter expressed from the *hsp-16.2* promoter^14^ showed strong induction throughout the worm body (**Fig. 2d and Extended Data Fig. 4a,b**). Exposing only part of the worm to heat shock resulted in fluorescence accumulation only in the same part, and to a lesser degree (**Fig 2e-f and Extended Data Fig. 4a,c-d and Movie 5)**. Thus, *hsp-16.2* is activated only in those parts of the worm that show DAF-16 nuclear accumulation.

Based on the observations that DAF-16 activation in body cells is preceded by activation in head neurons, we hypothesized that some head neurons are essential for DAF-16 activation by heat. We first examined three pairs of head sensory neurons (AFD, AWC and ASI) that have been implicated in thermosensation in the context of thermotaxis^15–18^. We profiled the activation of DAF-16 in genetic backgrounds that render each of these pairs inactive. Genetic ablation of AFD^19^ and AWC^20^ had only minor effect on the dynamics of activation (**Fig. 3a-c**). Inactivation of the ASI neurons^21^ — known to increase longevity in a DAF-16 dependent way^22^ — had little impact on the dynamics of activation, but displayed a prolonged recovery period (**Fig. 3a,d**). We thus concluded that none of the identified head thermosensory neurons are essential for DAF-16 heat-shock response.

**Figure 3.**
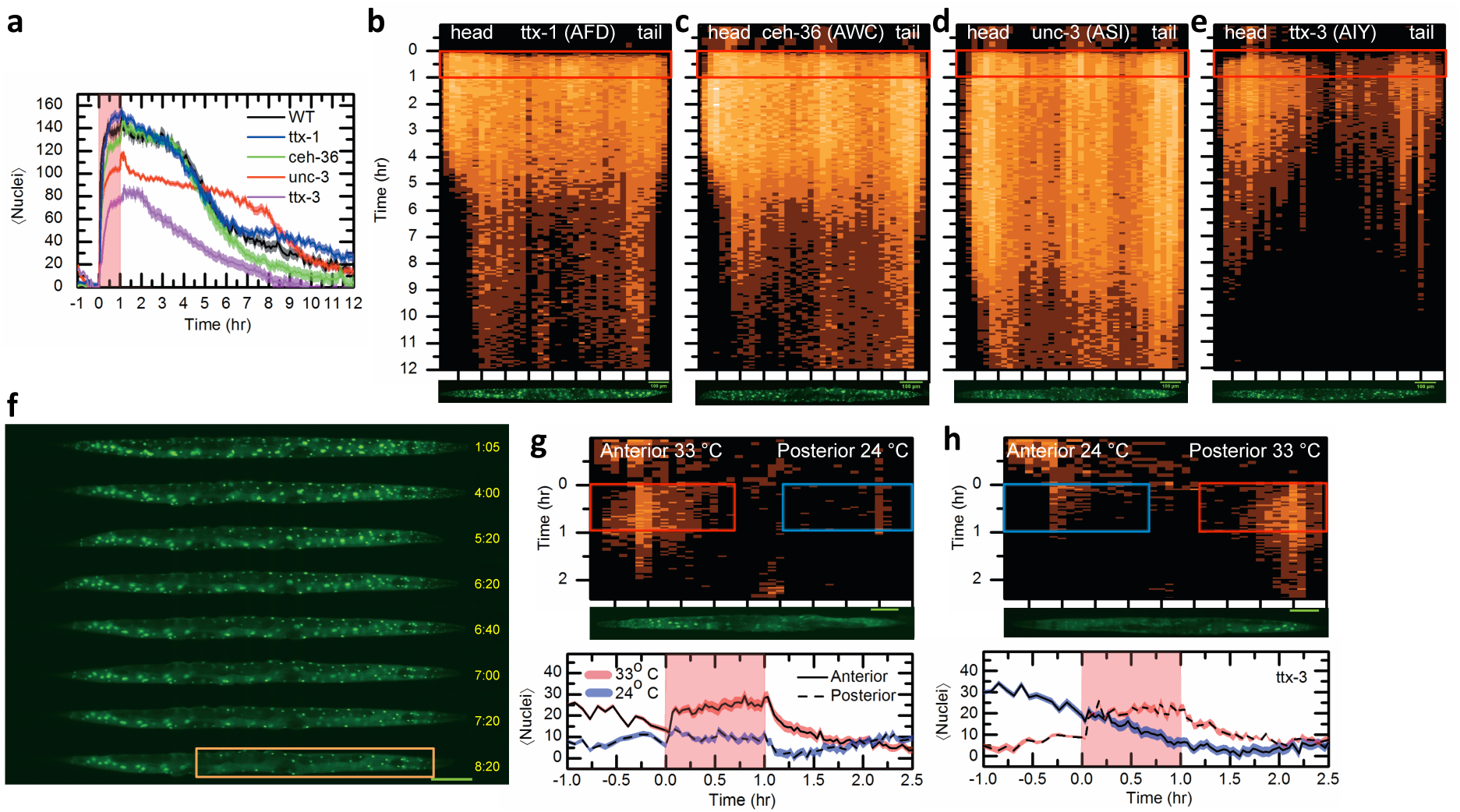
Effect of thermotactic neurons on DAF-16 heat shock response. **a,** Mean number of nuclei with localized fluorescence in the wild-type (N2) worms (black) and mutants carrying null alleles of *ttx-1* (ablating the ADF neurons, blue); *ceh-36* (AWC, green); *unc-3* (ASI, red); and *ttx-3* (AIY, purple). In all experiments a 33°C heat shock was applied at time zero and lasted one hour (pink shade). **b-e,** DAF-16 activation dynamics, as in Fig.1, in the same genetic backgrounds. **b,** *ttx-1,* **c,** *ceh-36*, **d,** *unc-3,* **e,** *ttx-3*. **f**, Recovery pattern in *ttx-3* worms differ significantly from wild-type, cf. Fig. 1h. Recovery starts in the middle of the worm, rather than in the head, and intestinal cells recover before muscle and epidermal cells (orange box). **g,** DAF-16 activation dynamics in *ttx-3* worms under anterior heat-shock, as in Fig. 2b. **h,** posterior heat-shock, as in Fig. 2c.

The interneurons AIY are known to be essential for thermotaxis^15^ and – together with the thermosensory neurons AFD – are required for activation of HSF-1 dependent heat-shock response^23^. Inactivation of AIY by a null allele of *ttx-3*^24^ resulted in severe reduction in DAF-16 activation. Under a uniform heat shock, DAF-16 accumulation was observed only in head and tail neurons, in the intestine, and in a limited number of body cells (**Fig. 3a,e**). In addition, the temporal order of tissue activation was significantly different from that of wild-type worms (**Fig. 3f**), showing a different ordering of tissue activation and losing the clear head-to-tail progression observed in wild-type worms. This strong effect of AIY ablation on DAF-16 activation dynamics in non-neuronal cells supports the idea that these dynamics are centrally controlled. Delivery of the heat shock to the anterior part of *ttx-3* worms resulted mostly in activation of head neurons and intestinal cells (**Fig. 3g**), while heat shock in the posterior part yield in addition a simultaneous activation of some posterior cells (**Fig. 3h**). Together, we conclude that the AIY neurons are essential for coordinated systemic activation of DAF-16 by heat shock.

While no thermosensory neuron proved essential for DAF-16 temperature response, we asked if inactivation of groups of sensory neurons can mitigate this response. Multiple sensory neurons in the head and tail express a cGMP-gated ion channel encoded by the genes *tax-2* and *tax-4*^25^. Mutation in these genes abolishes thermotactic behavior^26^. We found that the DAF-16 response to heat shock in *tax-4* worms was strongly suppressed throughout the worm, displaying a random pattern of activation in all but head neurons and intestinal cells (**Fig. 4a,b**), with a clear bias towards the anterior part of the worm even under uniform heat shock. Another group of sensory neurons rely on the TRPV channel expressed by the genes *ocr-2* and *osm-9*. Consistent with the known role of TRPV channels in temperature sensation in vertebrates and flies^27^, the OCR-2/OSM-9 channel is involved in determining the threshold for heat avoidance in *C. elegans*^28^. Accordingly, we found that mutation in *ocr-2* abolished all DAF-16 activation, except in intestinal cells (**Fig. 4a,c**). Taken together, these data suggest a role for sensory neurons in activation of DAF-16 under heat-shock.

**Figure 4.**
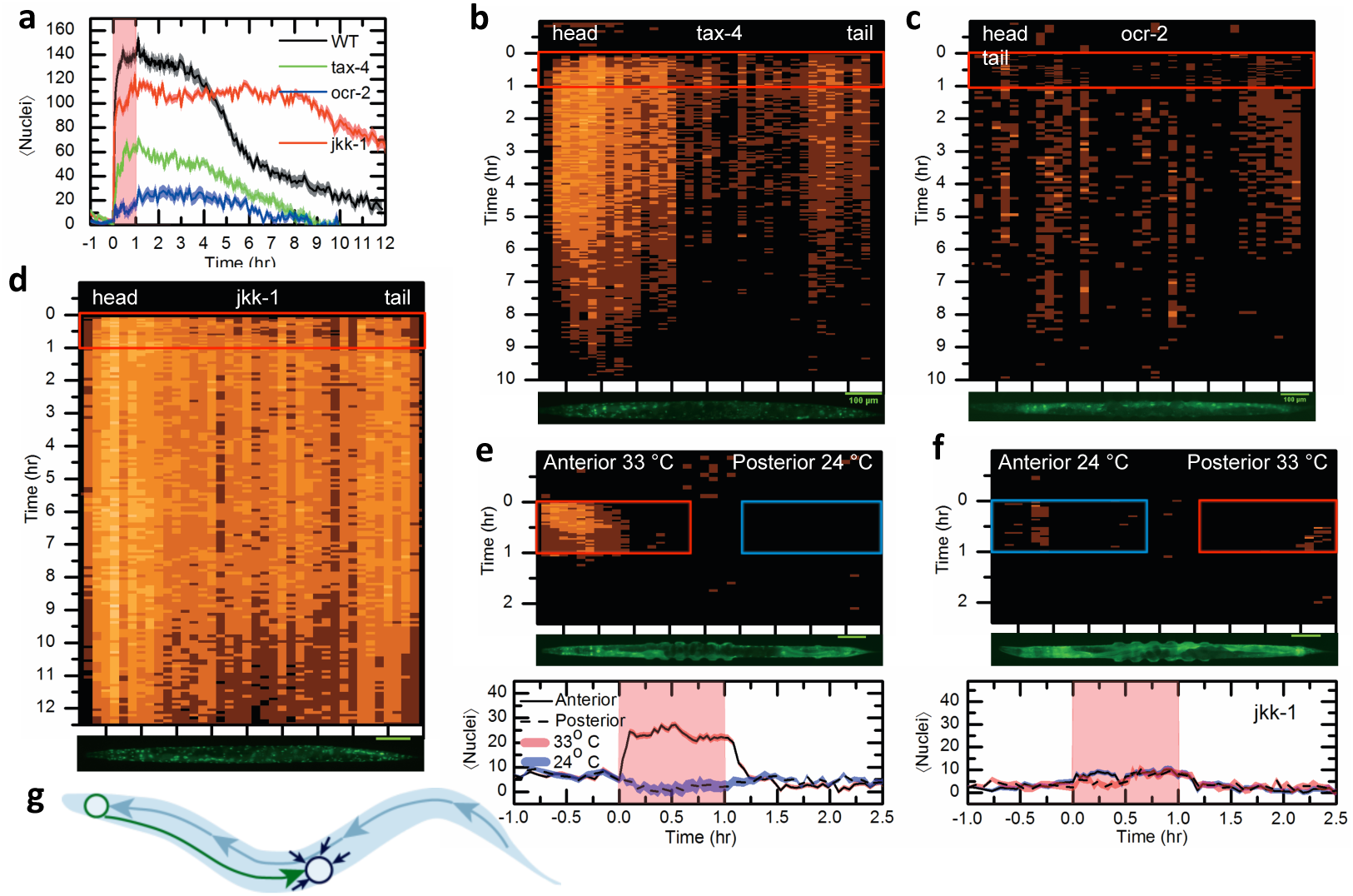
Sensory neurons and signal transduction are necessary for DAF-16 heat-shock response. **a,** Mean number of nuclei with localized fluorescence in the wild-type (N2) worms (black) and mutants carrying null alleles of *tax-4* (green); *ocr-2* (blue) and *jkk-1* (red).). In all experiments a 33°C heat shock was applied at time zero and lasted one hour (pink shade). **b-d,** DAF-16 activation dynamics, as in Fig.1, in the same genetic backgrounds: **b,** *tax-4*, **c,** *ocr-2* **d,** *jkk-1*. **e,** DAF-16 activation dynamics in *jkk-1* worms under anterior heat-shock, as in Fig. 2b. **f,** posterior heat-shock, as in Fig. 2c. **g,** Model. Temperature-related information collected throughout the worm body is transmitted to head sensory neurons (blue arrows) and integrated in interneurons, including AIY (blue circle). The integrated information is communicated to somatic cells from head to tail (green arrow). Activation of DAF-16 in response to heat shock in downstream cells (blue circle) requires this global signal, as well as local exposure to heat shock (black arrows) that exceeds a tissue-specific temperature threshold.

Our results suggest the existence of a signal transduction from head neurons to somatic tissues to activate DAF-16 in cells that experience elevated temperatures. One possible signaling mechanism is the JNK mitogen-activated protein kinase (MAPK) pathway. The activated MAPK JNK-1 is able to directly phosphorylate DAF-16 in neuronal cells^29^, and to promote nuclear translocation in non-neuronal cells in response to heat-shock^8^. In line with previous reports^29^, we find that inactivation of the JNK-1 pathway by mutation in the MAPK *jkk-1* has limited effect on DAF-16 activation by elevated temperature but showed difficulties in initiating DAF-16 deactivation (**Fig. 4d**). Exposure of the anterior part of the worm to elevated temperature resulted in activation of head neurons alone (**Fig. 4e**), while a heat shock focused on the posterior part of the worm resulted in no response at all (**Fig. 4f**). These findings suggest that the JNK pathway plays a role in information transfer leading to DAF-16 activation and deactivation, yet its precise functionality merits further investigation.

In summary, our results suggest a model (**Fig. 4g**) where DAF-16 activation occurs in cells exposed to a temperature that exceeds a tissue-specific temperature threshold, and requires a systemic signal that reflects the extent of exposure to high temperatures. Sensory neurons, in particular neurons that express a TRPV channel, and the JNK MAPK pathway are implicated in this process. Thus, DAF-16 response to elevated temperatures requires integration of local and systemic signals, ensuring a robust and coordinated response to molecular stress. These findings provide a framework for understanding the link between proteotoxicity and insulin in diabetes and aging^4,30^.

## Acknowledgements

We thank Ji Ying Sze for reagents, Yun Zhang, Mei Zhen and Aravi Samuel for discussions, Erin Dahlstrom and Lucy Eunju Lee for assistance, John Clarco and Susan Mango for critical reading of the manuscript. This work was supported in part by the National Science Foundations through grant PHY-1205494.

## Author Contributions

RBK and EL designed the research, analyzed the data and wrote the paper. RBK, KW and EL performed the experiments.

## Author Information

The authors declare no competing financial interests. Correspondence and requests for materials should be addressed to.

## Extended Data Figure Captions

**Extended Data Fig. 1.**
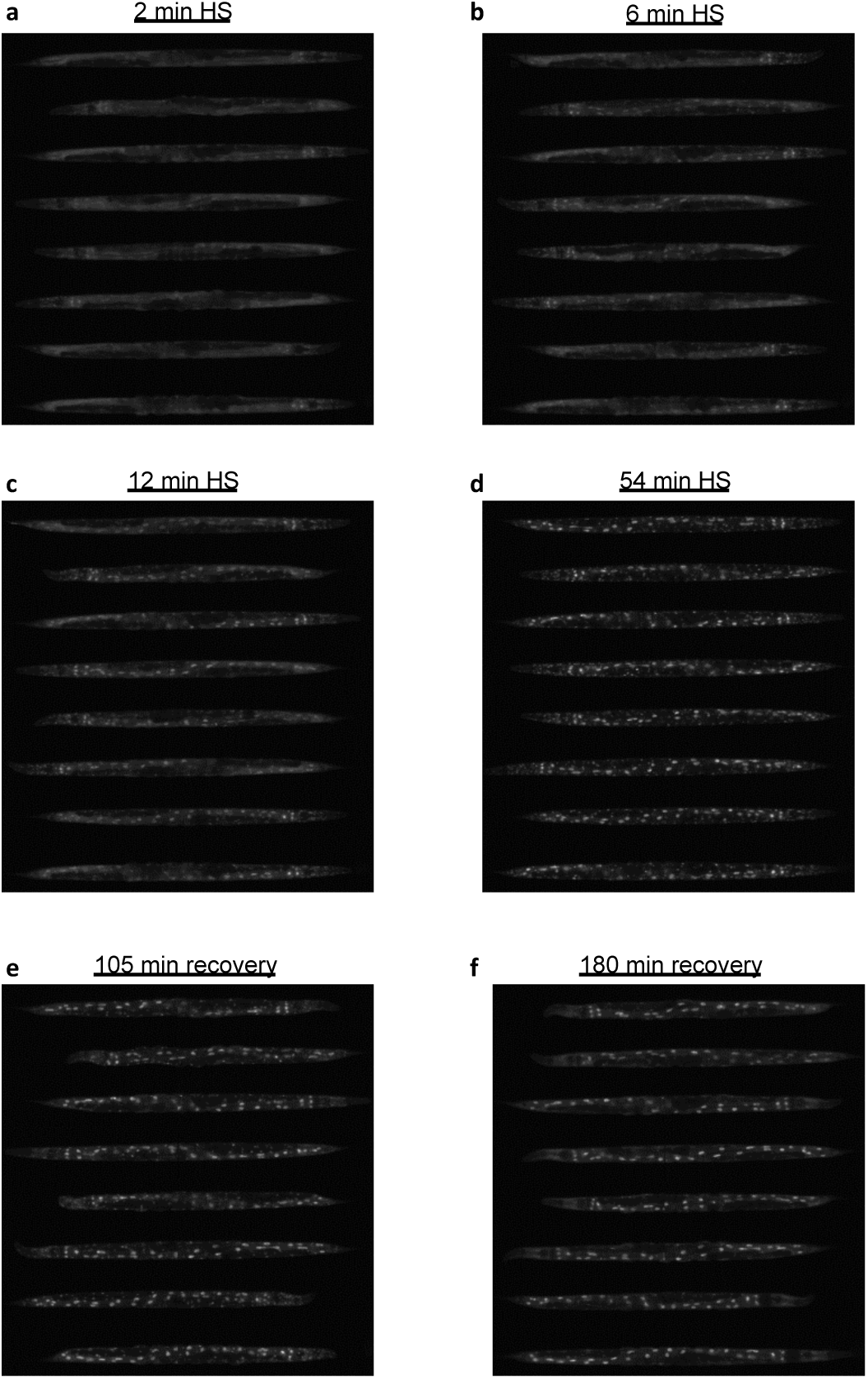
The dynamics of Daf-16 activation pattern is highly reproducible between worms. For demonstration, the localization pattern of 8 randomly selected worms is shown during (**a-d**) and after (**e-f**) a one-hour 32°C heat shock (HS).

**Extended Data Fig. 2.**
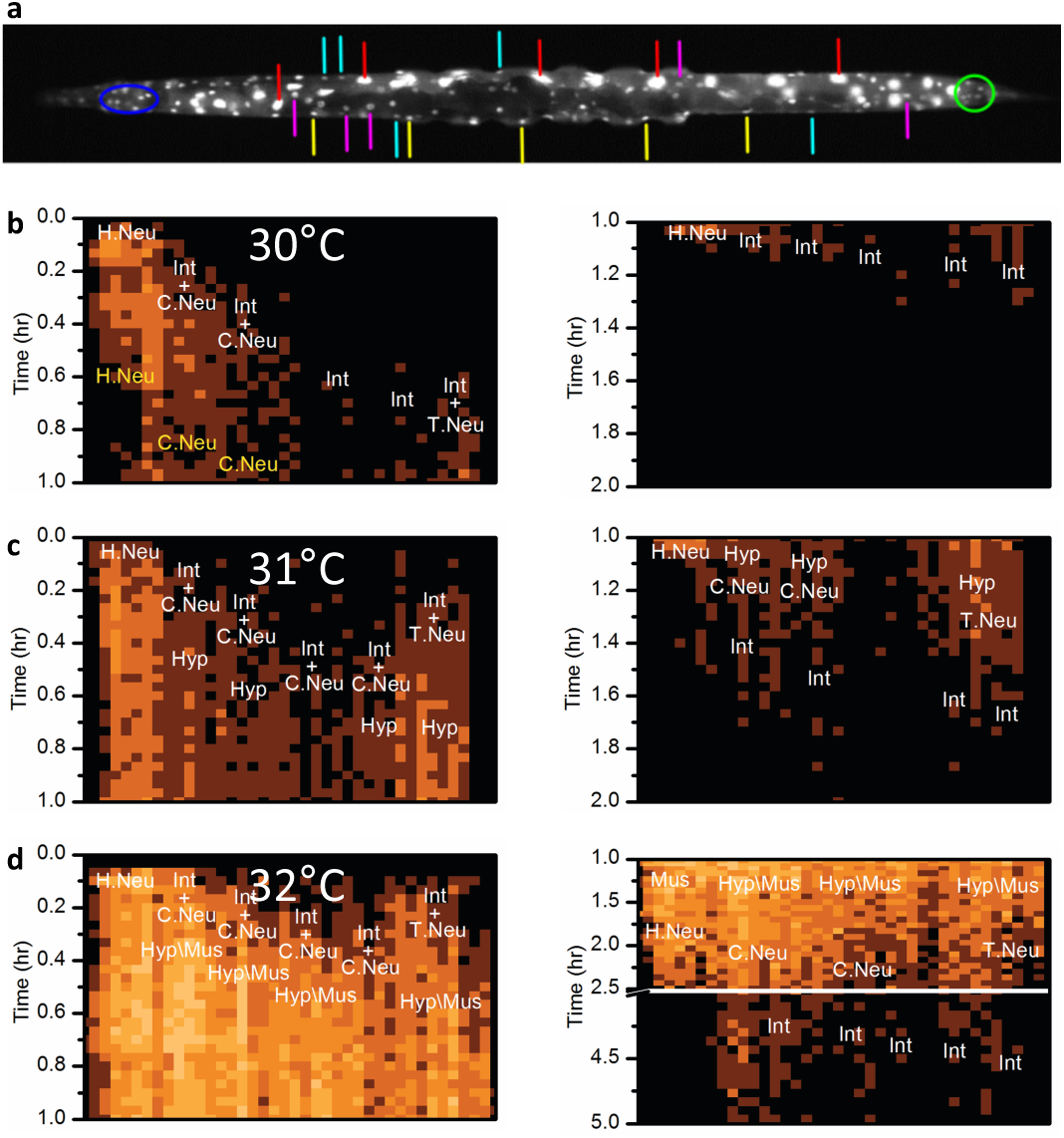
Tissue-specific DAF-16 dynamics in respond to heat shock. **a,** A representative subset of cells from different tissues was followed for determining tissue-specific timing of DAF-16 activation and deactivation: Intestinal cells (red), head neurons (circled in blue), tail neurons (circled in green), epidermal cells (purpule), ventral cord neurons (yellow), body-wall muscle cells (cyan). **b-d,** The spatiotemporal initiation of DAF-16 activation (left panel) and deactivation (right panel) during and after 30-32°C heat shocks. Cell types are marked as H.Neu - head neuron; T.Neu - Tail neuron; C.Neu – Ventral cord neuron; Int – Intestine; Hyp – Hypodermis; Mus – Muscle. Plus sign (+) represents simultaneous activation.

**Extended Data Fig. 3.**
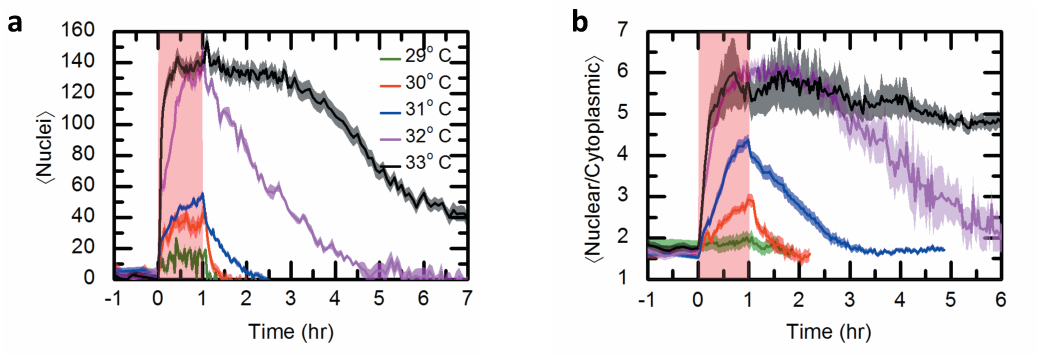
Heat shock response of DAF-16 is temperature dependent. **a,** Mean and standard error of the mean (s.e.m) of the number of localized nuclei as function of time. Worms were heat shocked for one hour (pink shade); time is presented in hours after the initiation of heat shock. **b,** Mean and s.e.m of the nuclear localization index, defined as the ratio between nuclear and cytoplasmic DAF-16 fluorescence.

**Extended Data Fig. 4.**
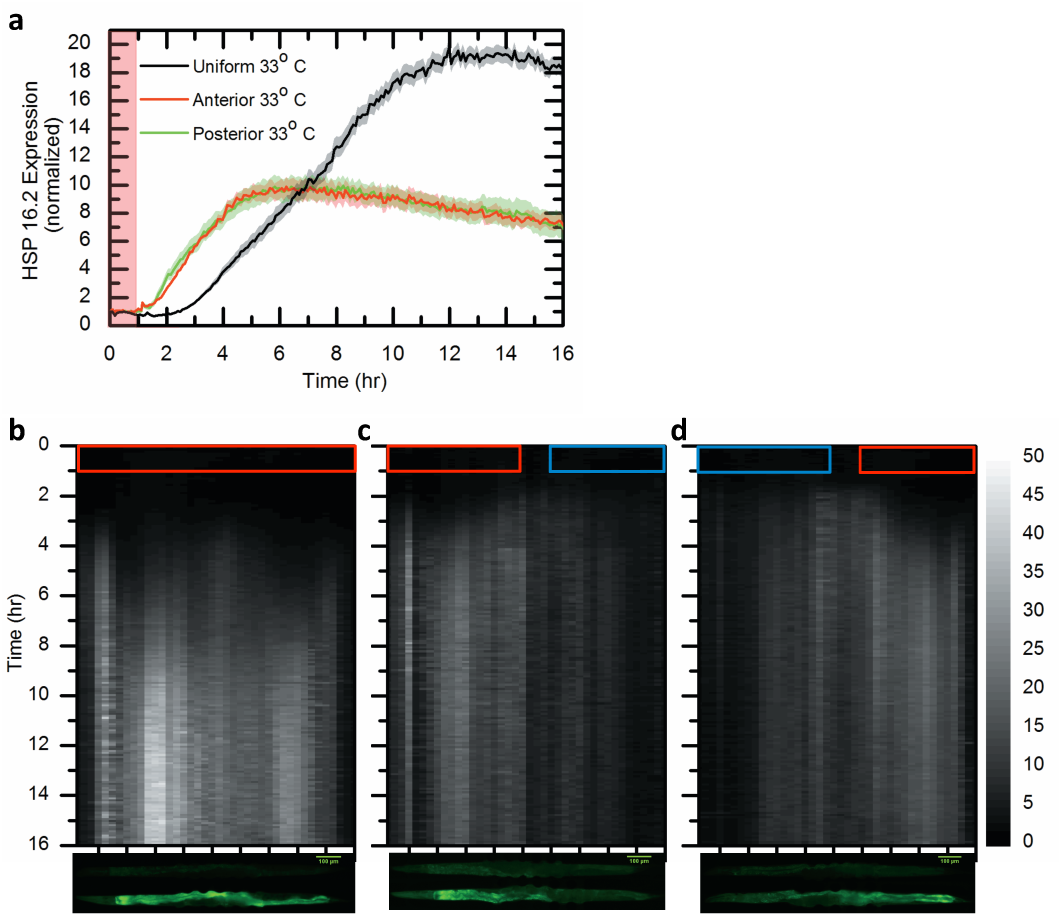
HSP16.2 expression in response to Heat shock. **a,** The mean and standard error of the mean (s.e.m) of total HSP-16.2::GFP fluorescence, normalized to total fluorescence before heat shock (HS). Black line corresponds to a uniform HS over the worm body, red and green lines to HS at the anterior or posterior parts, respectively. Worms were heat shocked for one hour (pink shade), time presented in hours after the initiation of heat shock. **b-d,** Mean spatiotemporal profile of HSP-16.2::GFP fluorescence, averaged over >30 worms, during and after a uniform (**b**), anterior (**c**) or posterior (**d**) heat shock (one hour, 32°C). Red boxes mark the heat-shock regions and times.

## Movie Captions

**Movie 1 | DAF-16 response to a 30°C heat shock.** DAF-16::GFP nuclear accumulation in response to a one hour heat shock at 30°C. Blue and red stripes indicate periods of ambient and heat-shock temperatures, respectively. Images were taken every 2 minutes. Time indicated in hours:minutes, scale bar 100um.

**Movie 2 | DAF-16 response to a 32°C heat shock.** DAF-16::GFP nuclear accumulation in response to a one hour heat shock at 32°C. Blue and red stripes indicate periods of ambient and heat-shock temperatures, respectively. Images were taken every 2 minutes. Time indicated in hours:minutes, scale bar 100um.

**Movie 3 | DAF-16 response to a 33°C heat shock.** DAF-16::GFP nuclear accumulation in response to a one hour heat shock at 33°C. Blue and red stripes indicate periods of ambient and heat-shock temperatures, respectively. Images were taken every 2 minutes. Time indicated in hours:minutes, scale bar 100um.

**Movie 4 | DAF-16 response to a spatial temperature step.** DAF-16::GFP nuclear accumulation in response to a one hour anterior or posterior heat shock. Blue and red stripes indicate ambient and heat-shock temperatures, respectively. Head of top worm is on the right, of bottom worm is on the left. Images were taken every 2 minutes during heat-shock, every 5 minutes before and every 4 minutes after. Time indicated in hours:minutes, scale bar 100um.

**Movie 5 | Induction of HSP-16 by a spatial temperature step.** Induction of HSP-16p::GFP in response to a one hour anterior or posterior heat shock. Blue and red stripes indicate ambient and heat-shock temperatures, respectively. Head of top worm is on the right, of bottom worm is on the left. Images were taken every 2 minutes during heat-shock and every 5 minutes before and after. Time indicated in hours:minutes, scale bar 100um.

## Methods

### Strains and worm preparation

*C. elegans* were maintained under standard conditions at 24C and fed *E. coli* OP50 bacteria. Wild-type worms were Bristol strain (N2). Other strains used in this study include: TJ356 zIs356 Is[Pdaf-16::daf-16-gfp; rol-6(su1006)]IV, TJ3001 zSi3001 [Phsp16.2::gfp::unc-54; cb-unc-119(+)]II, ERL25: ttx-1(p767) I; zIs356 IV, ERL26: tax-4(p678) III; zIs356 IV, ERL27: ttx-3(ks5) X; zIs356 IV, ERL31: jkk-1(km2) X; zIs356 IV, ERL42: ERL54: ceh-36(ks86) X; zIs356 IV, and ERL56: unc-3 (e151) X; zIs356 Worms were age-synchronized using standard protocol.

### Device fabrication and operation

The microfluidic device was made of two layers separated by a thin (100um) PDMS membrane. The first layer was used to accommodate worms, and the second layer for temperature control. Thickness of the first layer channels was 50 μm and of the second layer 300 μm. Food was delivered to the device using a computer actuated syringe pump (New-Era NE-501 OEM) actuated at flow rate of 5 μl/min. Short pulses (15 sec) of 150 μl/min were applied periodically (every 15min) to drive eggs into the egg-collection area.

### Video acquisition and data analysis

Imaging was done using a Zeiss Observer Z1 inverted microscope with 10X objective and a Hamamatsu Orca II camera with 50ms exposure at 7.8 pixels/10μm resolution. Images were taken every 2 minutes during the heat shock and every 5 minutes before and after. Image analysis was performed using custom MATLAB scripts, as detailed in the Supporting Methods.

